# Assigning enzyme sequences to orphan and novel reactions using knowledge of substrate reactive sites

**DOI:** 10.1101/210039

**Authors:** Noushin Hadadi, Homa MohamadiPeyhani, Ljubisa Miskovic, Marianne Seijo, Vassily Hatzimanikatis

## Abstract

Thousands of biochemical reactions with characterized activities are orphan, meaning they cannot be assigned to a specific enzyme, leaving gaps in metabolic pathways. Novel reactions predicted by pathway-generation tools also lack associated sequences, limiting protein engineering applications. Associating orphan and novel reactions with known biochemistry and suggesting enzymes to catalyze them is a daunting problem. We propose a new method, BridgIT, to identify candidate genes and protein sequences for these reactions, and this method introduces, for the first time, information about the *enzyme binding pocket* into reaction similarity comparisons. BridgIT assesses the similarity of two reactions, one orphan and one well-characterized, nonorphan reaction, using their substrate reactive sites, their surrounding structures, and the structures of the generated products to suggest protein sequences and genes that catalyze the most similar non-orphan reactions as candidates for also catalyzing the orphan ones.

We performed two large-scale validation studies to test BridgIT predictions against experimental biochemical evidence. For the 234 orphan reactions from KEGG 2011 (a comprehensive enzymatic reaction database) that became non-orphan in KEGG 2018, BridgIT predicted the exact or a highly related enzyme for 211 of them. Moreover, for 334 out of 379 novel reactions in 2014 that were later catalogued in KEGG 2018, BridgIT predicted the exact or highly similar enzyme sequences.

BridgIT requires knowledge about only three connecting bonds around the atoms of the reactive sites to correctly identify protein sequences for 93% of analyzed enzymatic reactions. Increasing to six connecting bonds allowed for the accurate identification of a sequence for nearly all known enzymatic reactions.

**SIGNIFICANCE STATEMENT:** Recent advances in synthetic biochemistry have resulted in a wealth of novel hypothetical enzymatic reactions that are not matched to protein-encoding genes, deeming them “orphan”. Nearly half of known metabolic enzymes are also orphan, leaving important gaps in metabolic network maps. Proposing genes for the catalysis of orphan reactions is critical for applications ranging from biotechnology to medicine. In this work, a novel computational method, BridgIT, identified a potential enzyme sequence to orphan reactions and nearly all theoretically possible biochemical transformations, providing candidate genes to catalyze these reactions to the research community. BridgIT online tool will allow researchers to fill the knowledge gaps in metabolic networks and will act as a starting point for designing novel enzymes to catalyze non-natural transformations.

## INTRODUCTION

Genome-scale reconstructions of metabolic networks can be used to correlate the genome with the observed physiology, though this hinges on the completeness and accuracy of the sequenced genome annotations. Orphan reactions, which are enzymatic reactions without protein sequences or genes associated with their functionality, are common and can be found in the genome-scale reconstructions of even well-characterized organisms, such as *Escherichia coli* (1). A recent review of orphan reactions reported that almost half of the enzymatic reactions cataloged in the Kyoto Encyclopedia of Genes and Genomes (KEGG) (2) lack an associated protein sequence (3).

Problems with orphan-like reactions can also arise in areas such as bioremediation, synthetic biology, and drug discovery, where exploring the potential of biological organisms beyond their natural capabilities has prompted the development of tools that can generate *de novo* hypothetical enzymatic reactions and pathways (4–14). These *de novo* reactions are behind many success stories in biotechnology, and they can also be used in the gap-filling of metabolic networks (5,11,12,14–17). While these enzymatic reactions have well-explained biochemistry that can conceivably occur in metabolism, they are essentially orphan reactions because they have no assigned enzyme or corresponding gene sequence. The lack of protein-encoding genes associated with the functionality of these *de novo* reactions limits their applicability for metabolic engineering, synthetic biology applications, and the gap-filling of genome scale models (18). A method for associating *de novo* reactions to similarly occurring natural enzymatic reactions would allow for the direct experimental implementation of the discovered novel reactions or assist in designing new proteins capable of catalyzing the proposed biotransformation.

Computational methods for identifying candidate genes of orphan reactions have mostly been developed based on protein sequence similarity (3,19–21). The two predominant classes of these sequence-based methods revolve around gene/genome analysis (21–24) and metabolic information (25, 26). Several bioinformatics methods combine different aspects of these two classes, such as gene clustering, gene co-expression, phylogenetic profiles, protein interaction data, and gene proximity, for assigning genes and protein sequences to orphan reactions (27–30). All of these methods use the concept of *sequence similarity* of the corresponding enzyme to determine the biochemical functions of orphan reactions.

This can be problematic because many known enzymatic activities are still missing an associated gene due to annotation errors, the incompleteness of gene sequences (31), and the fact that homology-based methods cannot annotate orphan protein sequences with no or little sequence similarity to known enzymes (3, 32). Moreover, sequence similarity methods can provide inaccurate results, as small changes in key residues might greatly alter enzyme functionality (33), and also it is a common observation that vastly different protein sequences can exhibit the same fold and, therefore, have similar catalytic activity even though they look very different (34, 35). In addition, these methods are not suitable for the annotation of *de novo* reactions since current pathway prediction tools only provide information about enzyme catalytic biotransformations and not about their sequences.

These shortcomings motivated the development of alternative computational methods based on the *structural similarity of reactants and products* for identifying candidate protein sequences for orphan enzymatic reactions (30,33,36–40). The idea behind these approaches was to assess the similarity of two enzymatic reactions via the similarity of their reaction fingerprints, i.e., the mathematical descriptors of the structural and topological properties of the participating metabolites (41), which could eliminate the problems associated with non-matching or unassigned protein sequences. In such methods, the reaction fingerprint of an orphan reaction is compared with a set of non-orphan reference-reaction fingerprints, and the genes of the most similar reference reactions are then assigned as promising candidate genes for the orphan reaction. Reaction fingerprints can be generated based on different similarity metrics, such as the bond change, reaction center, or structural similarity (40).

One class of reaction-fingerprint computational methods compares all of the compounds participating in reactions (40), which includes both reactants and cofactors. The application of this group of methods is restricted to specific enzymatic reactions that do not involve large cofactors (30,33,36–40). This is because the structural information of the large cofactors overwhelmingly contributes to the corresponding reconstructed reaction-fingerprint, and consequently, reactions with similar cofactors will inaccurately be classified as similar (35–38). Another class of reaction-fingerprint methods uses the chemical structures of reactant pairs for comparison (38). While these methods can be applied to all classes of enzymatic reactions, they neglect the crucial role of cofactors in the reaction mechanism. Moreover, neither of these two classes of methods have been employed for assigning protein sequences to *de novo* reactions (38).

In this study, we introduce a novel computational method, BridgIT, that links orphan reactions and *de novo* reactions, predicted by pathway design tools such as BNICE.ch (15), Retropath2 (14), DESHARKY (9), and SimPheny (11), with well-characterized enzymatic reactions and their associated genes. BridgIT uses reaction fingerprints to compare enzymatic reactions and is inspired by the “lock and key” principle that is used in protein docking methods (42) wherein the enzyme binding pocket is the “lock” and the ligand is a “key”. If a molecule has the same reactive sites and a similar surrounding structure as the native substrate of a given enzyme, it is then rational to expect that the enzyme will catalyze the same biotransformation on this molecule. Following this reasoning, BridgIT uses the structural similarity of the reactive sites of participating substrates together with their surrounding structure as a metric for assessing the similarity of enzymatic reactions. BridgIT introduces an additional level of specificity into reaction fingerprints by capturing critical information about the enzyme binding pocket. More precisely, BridgIT is substrate-reactive-site centric, and its reaction fingerprints reflect the specificities of biochemical reaction mechanisms that arise from the type of enzymes catalyzing those reactions.

Through several studies, we demonstrated the effectiveness of utilizing the BridgIT fingerprints for mapping novel and orphan reactions to the known biochemistry. These reactions are mapped according to the enzyme commission (EC) (43) number, which is an existing numerical classification scheme for enzyme-based reactions. The EC number can classify enzymes at up to four levels, with a one-level classification being the most general and a four-level classification being the most specific, and these enzyme-based reactions are then represented by four numbers, one for each level, separated by periods (e.g. 1.1.1.11). We show that BridgIT is capable of correctly predicting enzymes with an identical third-level EC number, indicating a nearly identical type of enzymatic reaction, for 94% of orphan reactions from KEGG 2011 that became non-orphan in KEGG 2016. This result validates the consistency of the sequences predicted by BridgIT with the experimental observations, and it further suggests that BridgIT can provide enzyme sequences for catalyzing nearly all orphan reactions. We also studied how the size of the BridgIT fingerprint impacts the BridgIT predictions. We show that BridgIT correctly identifies protein sequences using fingerprints that describe the neighborhood up to six bonds away from the atoms of the reactive site. Strikingly, we also find that it is sufficient to use the information of only three bonds around the atoms of the reactive sites of substrates to accurately identify protein sequences for 93% of the analyzed reactions.

Finally, to indicate the power of this computational technique, we applied BridgIT to the study of all of the 137,000 novel reactions from the ATLAS of biochemistry, a database of all theoretically possible biochemical reactions (44), most of which have no current route to their synthesis or development. Using our technology, we provide candidate enzymes that can potentially catalyze the biotransformation of these reactions to the research community, which should provide a basis for the engineering and development of novel enzyme-catalyzed biotransformations.

## RESULTS AND DISCUSSION

### BridgIT method

The BridgIT workflow together with an example of its application on an orphan reaction is demonstrated in Fig. 1. BridgIT is organized into four main steps (see Methods for more details): 1) reactive site identification, 2) reaction fingerprint construction, 3) reaction similarity evaluation, and 4) scoring, ranking, and gene assignment. The inputs of the workflow are (i) an orphan or a novel reaction and (ii) the collection of BNICE.ch generalized enzyme reaction rules. These reaction rules assemble biochemical knowledge distilled from the biochemical reaction databases, and they are used to discover *de novo* enzymatic reactions as well as predict all possible pathways from known compounds to target molecules (15,44,45). Here, we used the generalized enzyme reaction rules to extract information about the reactive sites of substrates participating in an orphan or a novel reaction, and we integrated it into the BridgIT reaction fingerprints (Fig. 1, panels 1 and 2). We then compared the obtained BridgIT reaction fingerprints to the ones from the reference reaction database based on the Tanimoto similarity scores (Fig. 1, panel 3). A Tanimoto score near 0 designates reactions with no or low similarity, whereas a score near 1 designates reactions with high similarity. We used these scores to rank the assigned reactions from the reference reaction database, and we identified the enzymes associated with the highest-ranked reference reactions as candidates for catalyzing the analyzed orphan or novel reaction (Fig. 1, panel 4). In the next sections, we discuss the reconstructions and testing of the various components of BridgIT as well as the results of our main analyses. A web-tool of BridgIT can be consulted at http://lcsb-databases.epfl.ch/pathways/Bridgit upon subscription.

**Fig. 1.**
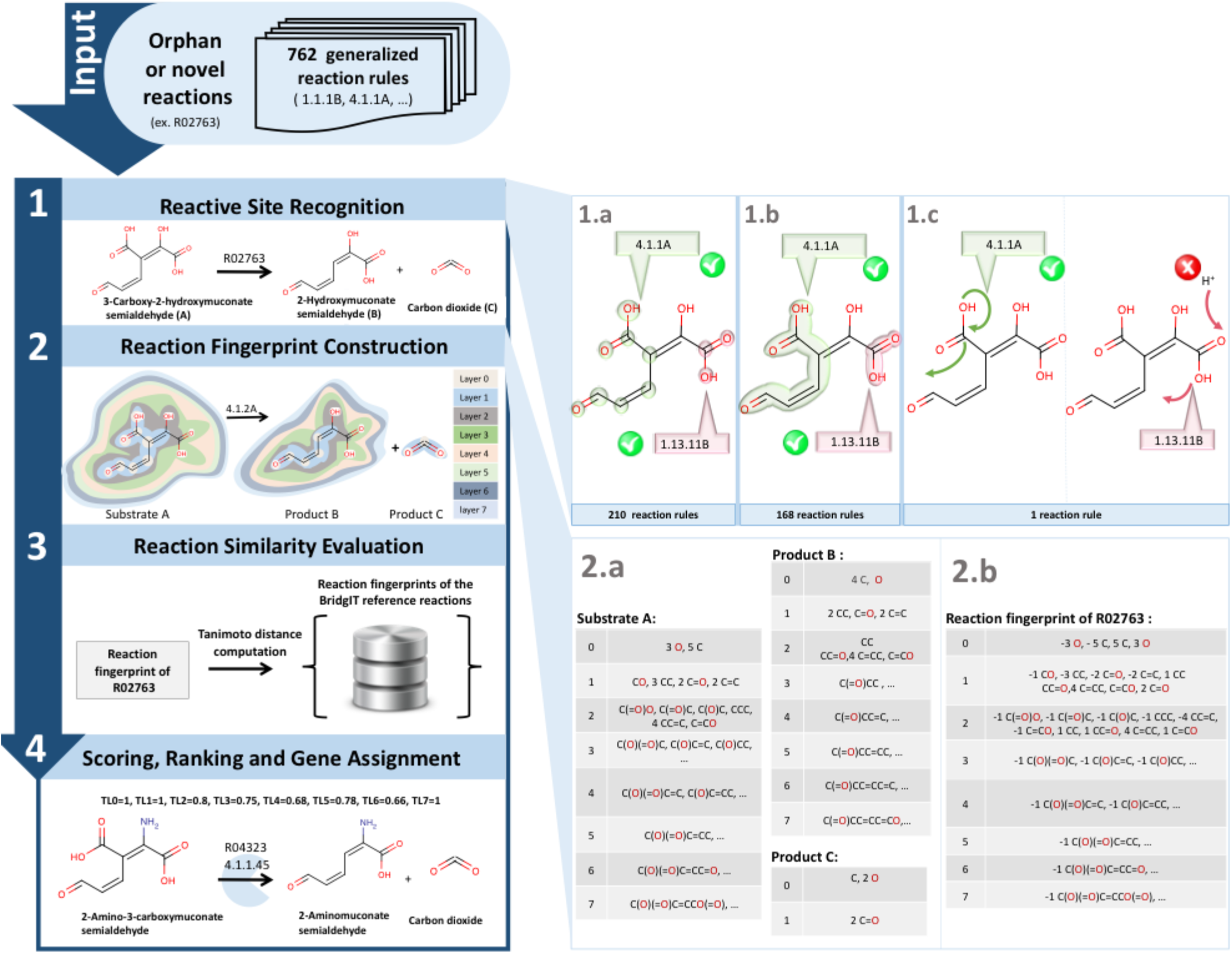
Main steps of the BridgIT workflow: (1) reactive site recognition for an input reaction *(de novo* or orphan); (2) reaction fingerprint construction; (3) reaction similarity evaluation; and (4) sorting, ranking and gene assignment. Panels 1.a to 1.c illustrate the procedure of the identification of reactive sites for the orphan reaction R02763. Panel 1.a: Two candidate reactive sites of 3-Carboxy-2-hydroxymuconate semialdehyde (substrate A) that were recognized by the rules 4.1.1. (green) and 1.13.11 (red). Panel 1.b: Both rules recognized the connectivity of atoms within two candidate reactive sites. Panel 1.c: Only reaction rule 4.1.1. can explain the transformation of substrate A to products. Panel 2.a shows the fragmentation of reaction compounds, whereas panel 2.b illustrates the mathematical representations of the corresponding BridgIT reaction fingerprints.

### Reference reaction database

The BridgIT reference reaction database is an essential component of the BridgIT workflow (Fig. 1). It consists of well-characterized reactions with associated genes and protein sequences, and it was built based on the KEGG 2016 reaction database (Methods). The KEGG database is the most comprehensive database of enzymatic reactions, and it provides information about biochemical reactions together with their corresponding enzymes and genes. However, half of KEGG reactions lack associated genes and protein sequences, and they are hence considered to be orphan reactions. The BridgIT reference database was built using the KEGG reactions that (i) can be reconstructed by the existing BNICE.ch generalized reaction rules and are elementally balanced (5,270 reactions) and (ii) are non-orphan (5,049 reactions). This restriction removes reactions that lack characterized substrate reactive sites, meaning that they cannot be used in our comparisons. As a result, the reference reaction database contains information for 5,049 out of 9,556 KEGG reactions (SI Dataset, Table S1).

### Sensitivity analysis of the BridgIT fingerprint size

The defining characteristic of the BridgIT reaction fingerprint is that it is centered around the reactive site of the reaction substrate(s). The number of description layers in the BridgIT fingerprint, i.e., the fingerprint size, defines how large of a chemical structure around the reactive site we consider when evaluating the similarity (Methods). To investigate to what extent the fingerprint size affects the similarity results, we performed a sensitivity analysis where we varied the fingerprint size between 0 to 10.

For this analysis, we considered the 5,049 non-orphan KEGG reactions that existed in the BridgIT reference reaction database. We started by forming reaction fingerprints that contained only the description layer 0 (fingerprint size 0) and evaluated how many of 5,049 nonorphan reactions BridgIT could correctly identify. We next formed the reaction fingerprints using only the description layers 0 and 1 (fingerprint size 1), and we performed the evaluation again. We repeated this procedure until the final step, where we formed the reaction fingerprints with ten description layers (fingerprint size 10).

As expected, the increase in the fingerprint size, i.e., specificity, led to a decrease in the average number of similar reactions assigned to the studied reactions. Moreover, the more description layers that were incorporated into the BridgIT fingerprint, the more accurately BridgIT matched the analyzed reactions (Table 1). Already for a fingerprint size 7, BridgIT correctly mapped 100% of the analyzed reactions, i.e., each of the 5,049 non-orphan reactions was matched to itself in the reference reaction database. This indicated that the information about six atoms along with their connecting bonds around the reactive sites was sufficient for BridgIT to correctly match all non-orphan KEGG reactions, and we chose the fingerprint size 7 for our further studies.

**Table 1.** Percent of correctly mapped reactions as a function of the size of the BridgIT fingerprint.

| Fingerprint size | 0 | 1 | 2 | 3 | 4 | 5 | 6 | 7 | 8 | 9 | 10 |
| --- | --- | --- | --- | --- | --- | --- | --- | --- | --- | --- | --- |
| % correctly mapped reactions | 4.3 | 35.2 | 60.5 | 72.1 | 92.7 | 97.8 | 98.6 | 100 | 100 | 100 | 100 |

### BridgIT reaction fingerprints offer improved predictions

We repeated the analysis from the previous section using the standard reaction difference fingerprint (Methods), which is used in structure similarity methods such as RxnSim (36) and RxnFinder (37), to assess the benefits of introducing the information about the reactive site of substrates into the reaction fingerprints. A comparison of the two sets of predictions on 5,049 non-orphan reactions showed that the predictions obtained with BridgIT-modified fingerprints were significantly better than the standard ones. BridgIT identified 100% of non-orphan reactions correctly versus the 71% success rate for the standard fingerprint method. Furthermore, BridgIT correctly matched 93% of the analyzed enzymatic reactions using the information about only three connecting bonds around the atoms of the reactive sites (fingerprint size 4), which exceeds the 71% of matched reactions when using the standard reaction fingerprints (fingerprint size 7) (Table 1).

The inferior performance of the standard reaction fingerprint method arose from three main sources. First, fragments from the substrate and product sets were cancelled out upon algebraic summation inside the fingerprint description layers (Methods), in which description layers 0 and 1 define the single atoms and the connected pairs of atoms of the reactive site, and layers 2 to 7 include information about the chemical structure around the reactive site that contains up to eight atoms and seven bonds (Fig. 1). This cancellation occurred in all description layers (fingerprint size 7) for 246 non-orphan reactions, i.e., their standard fingerprints were empty (SI Dataset, Table S3). As an example, Fig. 3A shows the standard reaction fingerprint of KEGG reaction R00722 that was empty for the standard fingerprint method. The information about reactive sites introduced in the BridgIT reaction fingerprints prevents such cancellations, since BridgIT does not include the atoms of the reactive site(s) in the process of the algebraic summation of the substrate and product set fragments (Methods). As a result, BridgIT mapped R00722 to itself and identified R00330 as the most similar reaction to R00722 (Fig. 2A). Indeed, according to the KEGG database, the enzyme 2.7.4.6 catalyzes both reactions.

**Fig. 2.**
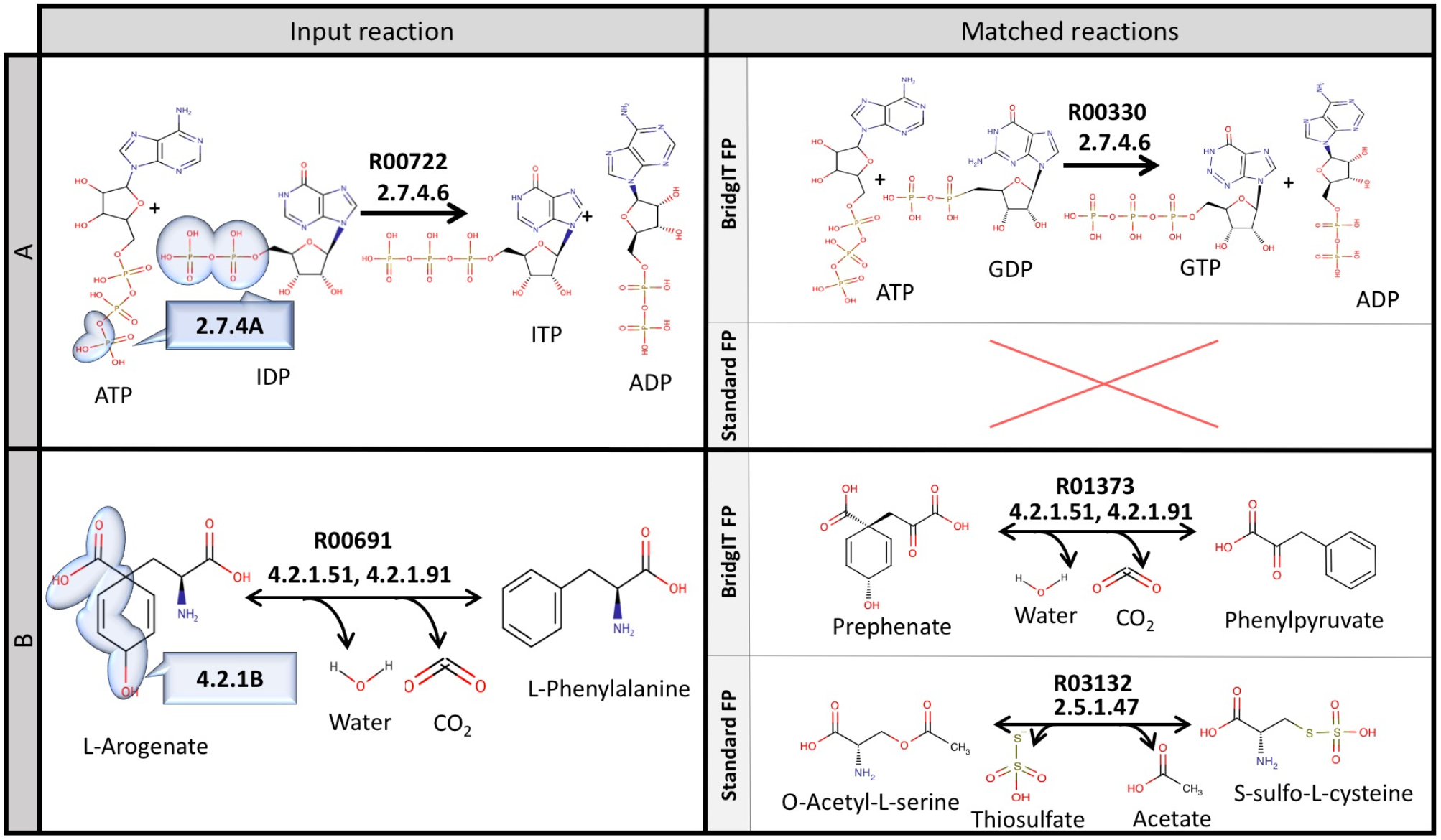
Comparison of the results obtained with the BridgIT and standard fingerprint on two example KEGG reactions. (A) The input reaction R00722 (left) and the most similar reactions (right) identified with the BridgIT and standard fingerprints. Note that the standard fingerprinting method failed to find a similar reaction to R00722 due to cancellations inside all fingerprint description layers. (B) The input reaction R00691 (left) and the most similar reactions (right) identified with the BridgIT and standard fingerprints.

**Fig. 3.**
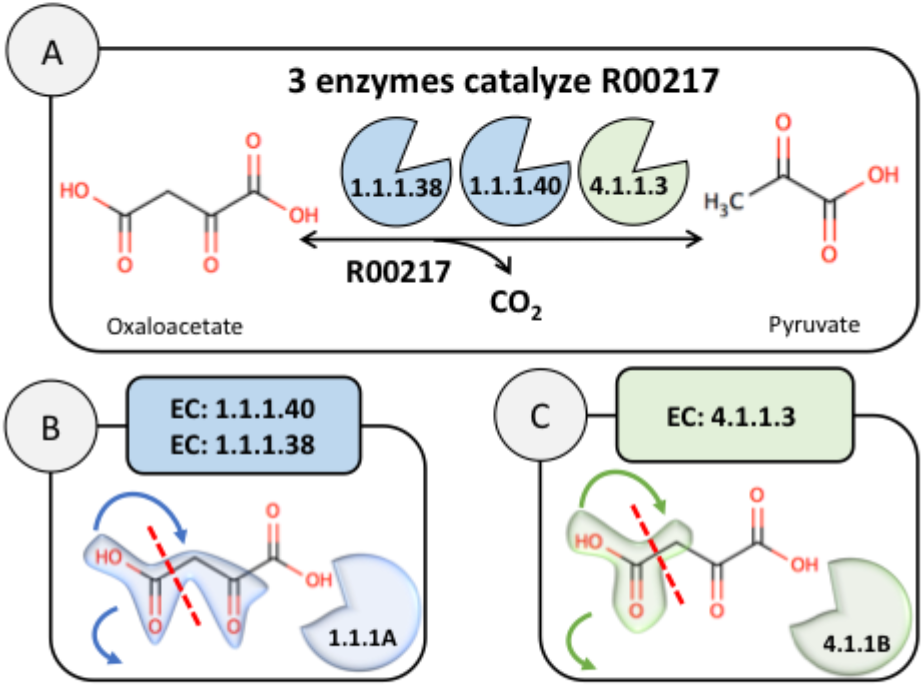
A multi-enzyme reaction such as R00217 can be catalyzed by more than one enzyme. BridgIT identified two distinct fingerprints for this reaction that correspond to two reactive sites of oxaloacetate. The reactive site recognized by the 1.1.1.-rule is more specific (blue substructure) than the one recognized by the 4.1.1.- rule (green substructure).

Second, the performance of the standard reaction fingerprint suffered because the first description layer of the standard fingerprint was empty for an additional 1,129 reactions, which indicated that these fingerprints did not represent the bond changes during the reaction (SI Dataset, Table S4).

Third, the remaining 89 mismatched non-orphan reactions had partial cancellations in the fingerprint description layers. For example, the standard fingerprint method incorrectly identified R03132 as the most similar to R00691, whereas BridgIT identified R00691 and R01373 as the most similar to R00691 (Fig. 2B), which matches the KEGG reports indicating that both R00691 and R01373 can be catalyzed by either EC 4.2.1.51 or EC 4.2.1.91.

### From reaction chemistry to detailed enzyme mechanisms

Approximately 15% of KEGG reactions (1,532 reactions) are assigned to more than one enzyme and EC number, i.e., multiple enzymes can catalyze a specific biotransformation through different enzymatic mechanisms. For example, KEGG reaction R00217 is assigned to three different EC numbers, 4.1.1.3 (oxaloacetate carboxy-lyase), 1.1.1.40, and 1.1.1.38 (both malate dehydrogenases), and the corresponding reactions involve different mechanisms (Fig. 3). For the 4.1.1.3 enzyme, the reaction mechanism is well understood, as this enzyme belongs to the carboxy-lyases, where a carbon-carbon bond is broken and a molecule of CO2 is released. In contrast, for the corresponding enzymes 1.1.1.40 and 1.1.1.38, there is ambiguity about their detailed mechanisms. As discussed in Swiss-Prot (46), these two enzymes are both NAD-dependent dehydrogenases that also have the ability to decarboxylate oxaloacetate. They are found in bacteria and insects (1.1.1.38) or in fungi, animals, and plants (1.1.1.40). In addition, BNICE.ch identifies multiple alternative reactive sites for 42% of the KEGG reactions that have a single enzyme assigned to them. Consequently, multiple reaction fingerprints can describe the biotransformation of these reactions.

We applied the BridgIT algorithm to R00217 in order to see how well BridgIT matched this reaction to its known enzymes, and we obtained two distinct reaction fingerprints that corresponded to the two different enzyme mechanisms mentioned above. More precisely, the BNICE.ch generalized reaction rules 1.1.1.- and 4.1.1.- identified two different reactive sites of oxaloacetate to break the carbon-carbon bond and release CO2 and pyruvate (Fig. 3). The 1.1.1.- rule recognized a larger, i.e., more specific, reactive site compared to the one recognized by 4.1.1.- (Fig. 3).

Therefore, a single reaction from KEGG was translated into more than one fingerprint in the BridgIT reference database. This way, by preserving the information about enzyme binding pockets, the reconstructed BridgIT reference reaction database expands from 5,049 *reactions* to 17,657 *reaction fingerprints* corresponding to 17,657 *detailed reaction mechanisms.*

### Comparison of BridgIT and BLAST predictions

As a means to relate *reaction structural similarity* obtained using BridgIT with *reaction sequence similarity* obtained using BLAST (47), we applied these two techniques in parallel on a subset of reactions and their corresponding protein sequences from the reference reaction database. We compared the similarity results of BridgIT with those of BLAST, and we statistically assessed BridgIT performance using receiver operating characteristic (ROC) curve analysis (SI Fig. 1).

We chose *E. coli* BW29521 (EBW) as our benchmark organism for this analysis. We extracted all of the nonorphan reactions of EBW from the BridgIT reference database together with their associated protein sequences (SI Dataset, Table S2). There were 531 nonorphan reactions in EBW associated with 413 protein sequences. In total, there were 731 reaction-gene associations (SI Dataset, Table S2), as there were reactions with more than one associated gene and genes associated with more than one reaction. We then used BridgIT to assess the structural similarity of the 531 EBW reactions to the BridgIT reference reactions using the Tanimoto score, and we also applied BLAST to quantify the similarity of the 413 EBW protein sequences to the KEGG protein sequence database using e-values. We provided a list of BridgIT reaction-reaction comparisons together with BLAST sequence-sequence comparisons (SI Dataset, Table S2).

*Comparing reaction (BridgIT) and sequence (BLAST) similarity scores.* We considered two sequences to be similar if BLAST reported an e-value of less than 10^−10^ for their alignment. For a chosen discrimination threshold (DT) of the global Tanimoto score (T_G_) we considered the BridgIT prediction of similarity between an EBW reaction and a BridgIT reference reaction with a Tanimoto score of T_G_ as:

i. True Positive (TP) if T_G_ > DT and their associated sequence(s) were similar (e-value < 10^−10^);
ii. True Negative (TN) if not similar for both BridgIT (T_G_ < DT) and BLAST+ (e-value > 10^−10^);
iii. False Positive (FP) if similar for BridgIT (T_G_ > DT) but not similar for BLAST+ (e-value > 10^−10^);
iv. False Negative (FN) if not similar for BridgIT (T_G_ < DT) but similar for BLAST+ (e-value < 10^−10^).

We then counted the number of TPs, TNs, FPs, and FNs for all 531 reactions, and we summed these quantities to obtain the total number of TPs, TNs, FPs, and FNs per chosen DT. We repeated this procedure for a set of DT values varying across the interval between 0 and 1. Finally, we used the total number of TPs, TNs, FPs, and FNs to compute the true positive and false positive rates for the ROC curve analysis (SI Fig. 1A). The ROC curve indicated that the reaction comparison based on *reaction structural similarity* (BridgIT) was comparable to the one based on *reaction sequence similarity* (BLAST). Indeed, the obtained area under the ROC curve (AUC) score for the BridgIT classifier was 0.91, indicating that the similarities between the two methods were very high (SI Fig. 1A). We next studied if the type of compared reactions affected the accuracy of BridgIT predictions by categorizing reactions according to their first-level EC class, which indicates the broadest category of enzyme functionality, and then performing the ROC analysis for each class separately (SI Fig. 1A). The analysis revealed that BridgIT performed well with all major enzyme classes, as represented by the high AUC scores, ranging from 0.88 (EC 1) to 0.96 (EC 5).

We next analyzed the accuracy of BridgIT classification as a function of the DT of the Tanimoto score (SI Fig. 1B). The accuracy ranged from 43% for DT = 0.01 to 85% for DT = 0.30. For values of DT > 0.30, the accuracy monotonically decreased toward a value of 62% for DT = 1. The classifier was overly conservative for values of DT > 0.30, and it was rejecting true positives (SI Fig. 1B). More specifically, for DT = 0.30, the TP percentage was 38%, whereas, for DT = 1, it was reduced to 3%. In contrast, the TN percentage increased very slightly for the values of DT > 0.30, where for DT = 0.30, it was 46%, and for DT = 1, it was 57% (SI Fig. 1B). Based on this analysis, we have chosen a DT of 0.30 as an optimal threshold value for further studies.

### BridgIT analysis of known reactions with common enzymes

The 5,049 reactions in the reference database were catalyzed by only 2,983 enzymes, i.e., there were promiscuous enzymes that catalyzed more than one reaction. Out of the 2,983 enzymes, 844 of them were promiscuous, catalyzing 2,432 of the reactions (SI Dataset, Table S5). Interestingly, BridgIT correctly assigned more than 80% of these 2,432 reactions to their corresponding promiscuous enzyme. An example of such a group is given in Table 2. This table shows the same enzymes listed across the top and down the size of the grid, with the corresponding Tanimoto scores indicating the accuracy of BridgIT’s classifications. The overall high scores in this grid indicate the accuracy of the enzyme assignments.

**Table 2.** A group of five reactions catalyzed by enzyme 1.1.1.219, wherein the Tanimoto score is given for the comparison between the reaction listed across the top and the reaction listed down the side.

| Catalyzed reactions | R03123 | R03636 | R05038 | R07999 | R07998 |
| --- | --- | --- | --- | --- | --- |
| R03123 | 1 | 0.96 | 0.93 | 0.93 | 0.98 |
| R03636 | 0.96 | 1 | 0.96 | 0.94 | 0.95 |
| R05038 | 0.93 | 0.96 | 1 | 0.97 | 0.91 |
| R07999 | 0.93 | 0.94 | 0.97 | 1 | 0.91 |
| R07998 | 0.98 | 0.95 | 0.91 | 0.91 | 1 |

We investigated the remaining 20% of reactions in depth, and we observed that the Tanimoto scores of the first two description layers (Methods) indicated a very low similarity between the reactions catalyzed by the same enzyme. This result suggested that such enzymes were either multi-functional, i.e., they had more than one reactive site (Fig. 4), or were incorrectly classified in the EC classification system.

**Fig. 4.**
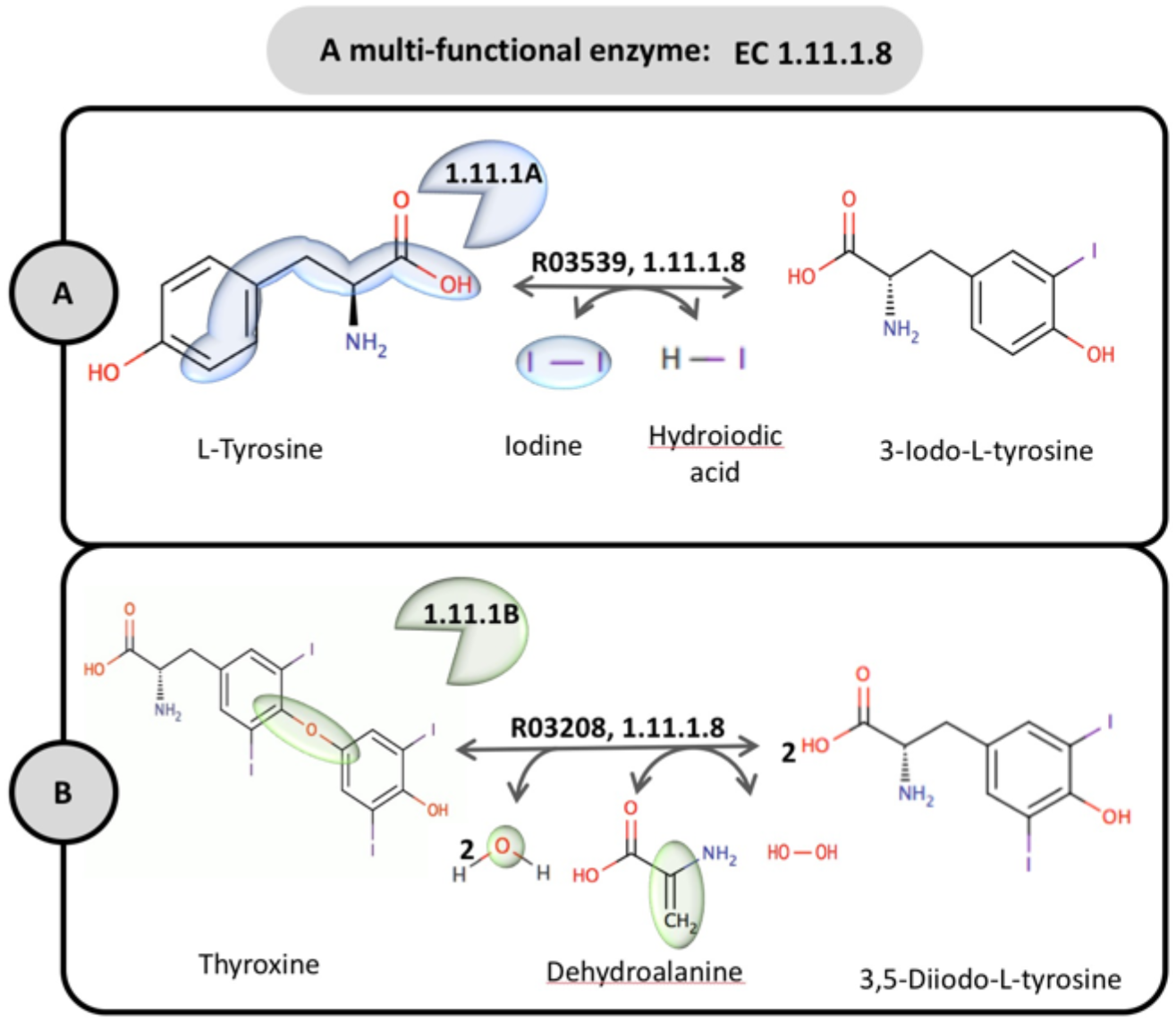
Multi-functional enzymes can catalyze reactions with two different reactive sites. (A) R03539 and (B) R03208 are catalyzed by the same enzyme, 1.11.1.8. However, the reactive sites of these substrates are completely different.

### BridgIT validation against biochemical assays

To assess BridgIT’s performance using biochemically confirmed reactions, we performed two validation studies on sets of (I) orphan and (II) novel reactions. Since the known reactions in KEGG are all experimentally confirmed using biochemical assays, we could use this pooled experimental data from hundreds of laboratories to demonstrate BridgIT’s ability to identify the activity of biologically relevant orphan reactions on a large scale.

*<u>Study I</u>:* We compared the number of orphan reactions in the two versions of the KEGG reaction database, KEGG 2011 and KEGG 2018. We found that 234 orphan reactions from KEGG 2011 were later associated with enzymes in KEGG 2018, meaning they became non-orphan reactions (SI Dataset, Tables S6-8). Since these newly classified reactions have been experimentally confirmed, we used these 234 reactions as a benchmark to evaluate BridgIT performance.

We formed the reference reaction database using the reactions from KEGG 2011 (Methods), and we compared the BridgIT results with the KEGG 2018 enzyme assignments up to the third EC level. Remarkably, BridgIT and KEGG 2018 assigned enzymes matched to the third EC level for 211 out of 234 (90%) reactions (SI Dataset, Tables S6 and S7). This means that BridgIT accurately predicted the enzyme mechanism and provided highly related protein sequences for enzymes that have been biochemically confirmed to catalyze a large majority of the orphan reactions in 2011.

The 234 reactions are catalyzed by 168 enzymes with specified fourth-level EC numbers in KEGG 2018. However, only 29 out of these 168 enzymes were cataloged in KEGG 2011, and the remaining 139 enzymes had new fourth-level EC classes assigned in KEGG 2018 – meaning BridgIT only had access to the 29 enzymes that were classified in KEGG 2011 from which the reference reaction database was built. The 29 enzymes catalyzed 35 out of the 234 studied reactions. For 29 out of these 35 (83%) orphan reactions, the BridgIT algorithm predicted the same sequences that KEGG 2018 assigned to these reactions (SI Dataset, Table S9). A higher matching score when comparing up to the third EC level rather than the fourth EC level is likely because BridgIT uses BNICE.ch generalized reaction rules, which describe the biotransformations of reactions with specificities up to the third EC level.

*<u>Study II</u>*: The ATLAS of biochemistry (44) provides a comprehensive catalog of theoretically possible biotransformations between KEGG compounds, and it can be mined for novel biosynthetic routes for a wide range of applications in metabolic engineering, synthetic biology, drug target identification, and bioremediation (40). We studied the 379 reactions from the ATLAS of Biochemistry that were novel in KEGG 2014 and were later experimentally identified and catalogued in KEGG 2018. We formed the reference reaction database using the reactions from KEGG 2014 and applied BridgIT to these 379 reactions. For 334 out of these 379 reactions, BridgIT proposed similar known reactions with a Tanimoto score higher than 0.3, thus providing promising protein sequences for enzymes catalyzing these reactions (SI Dataset, Table S10). For 14 of these novel reactions, BridgIT assigned the same sequences that were assigned in KEGG 2018 (SI Dataset, Table S11). An example of such a reaction is rat132341, which was a novel reaction in 2014 and later was catalogued as R10392 in KEGG 2018 (Fig. 5A). The BridgIT analysis of this reaction revealed that R03444, which is catalyzed by enzyme 4.2.1.114, is the structurally closest reaction to this novel one, suggesting that protein sequences from EC 4.2.1.114 can catalyze this novel reaction. This was later confirmed by experimental biochemical evidence, as R10392 is associated with the same EC 4.2.1.114 enzyme in KEGG 2018. There are 243 available protein sequences for enzyme 4.2.1.114, and one sequence already has a confirmed protein structure (Fig. 5C). This represents the first computational method for predicting protein sequences for orphan and novel reactions whose results were validated using experimental biochemical evidence on a large scale.

**Fig. 5.**
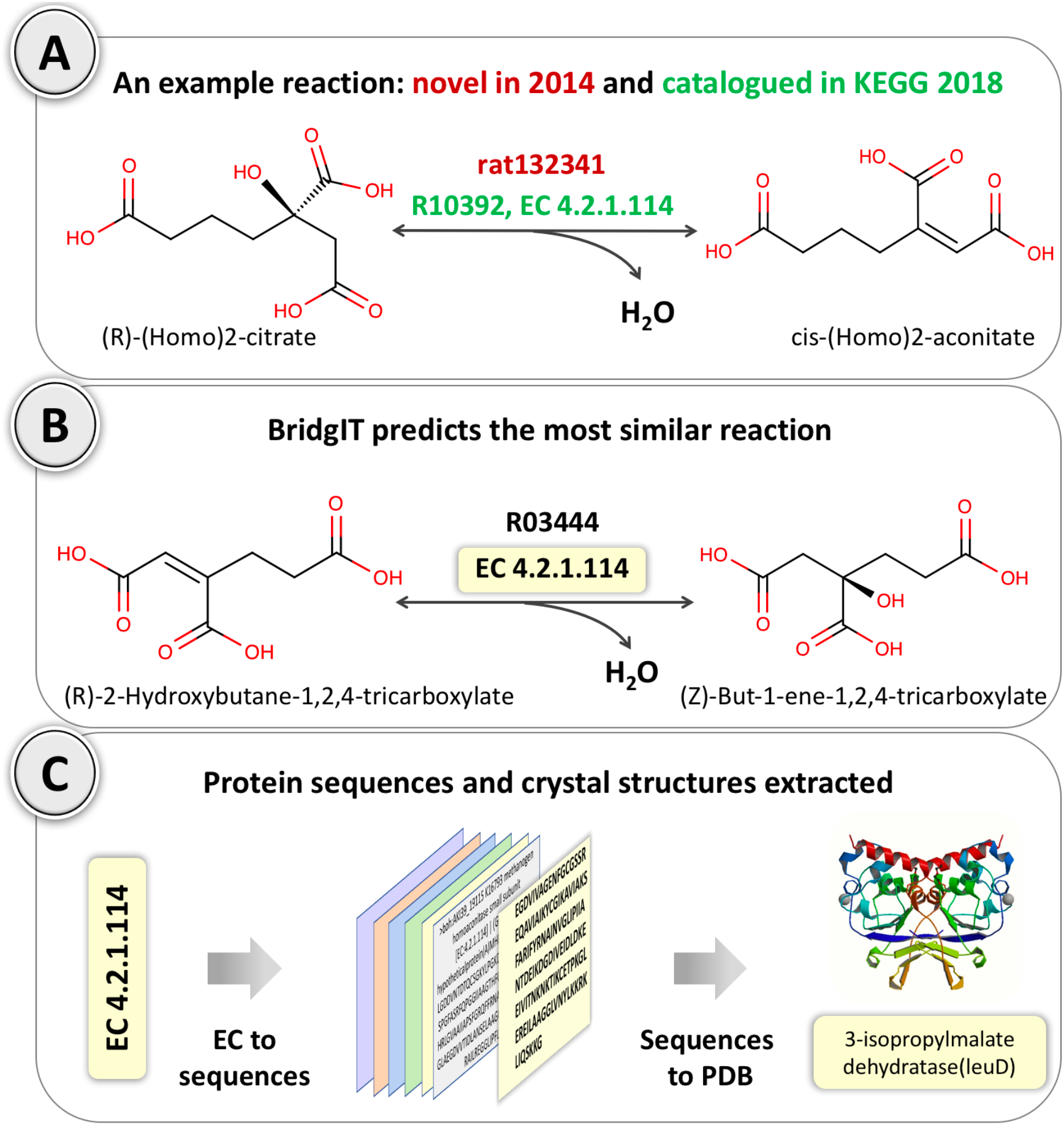
Details of the BridgIT verification procedure that was performed on ATLAS reaction rat132341, which was novel in KEGG 2014 and later experimentally identified and catalogued in KEGG 2018 — i.e., it became a non-orphan reaction (R10392). (A) rat132341 catalyzes the conversion of (R)-(Homo)2-citrate to cis-(Homo)2-aconitate. (B) Using the biochemical knowledge of KEGG 2014, BridgIT predicts the KEGG reaction R03444, which is catalyzed by a 4.2.1.114-class enzyme, as the most similar known reaction to rat132341. Remarkably, the same enzyme is later assigned to R10392 in KEGG 2018 with the corresponding biochemical confirmation. (C) The identified EC number (4.2.1.114) can be used to extract the corresponding protein sequences along with their crystal structures.

### BridgIT predictions for KEGG 2018 orphan reactions

We applied BridgIT to the 810 orphan KEGG 2018 reactions that could be reconstructed using the BNICE.ch generalized reaction rules. Remarkably, BridgIT identified corresponding reference reactions with Tanimoto scores higher than the optimal threshold value of 0.30 for 97% of the orphan reactions. The remaining 3% of orphan reactions had a low similarity with the reference reactions. This result and the fact that BridgIT correctly mapped 100% of non-orphan KEGG reactions suggested that, as our knowledge of biochemistry expands, the annotation of novel and orphan reactions using tools such as BridgIT will also improve.

### BridgIT predictions for ATLAS novel reactions

We further utilized BridgIT to identify candidate enzymes for all the 137,0 *de novo*, orphan-like, ATLAS reactions. These candidate enzymes can either be used directly in systems biology designs if the matched enzymes perform the desired catalysis, or their amino acid sequences can be optimized through protein engineering to achieve the desired results. We found that 7% of novel ATLAS reactions were matched to known KEGG reactions with a Tanimoto score of 1 (perfect match), while 88% were similar to KEGG reactions with a Tanimoto score higher than the optimal threshold value of 0.3. Therefore, BridgIT could identify promising enzyme sequences for catalyzing 95% of novel ATLAS reactions. The remaining 5% of these reactions were not similar to any of the well-characterized, known enzymatic reactions.

Finding well-characterized reactions that are similar to novel ones is crucial for evolutionary protein engineering as well as computational protein design, and methods like BridgIT can be instrumental in moving from a concept to the experimental implementation of *de novo* reactions. Additionally, to facilitate the experimental implementation of novel ATLAS reactions in metabolic engineering, systems and synthetic biology, and bioremediation studies, we can use the BridgIT similarity scores as confidence measures for evaluating the feasibility.

The results of the BridgIT analysis of the KEGG 2018 orphan and novel ATLAS reactions are available on the website http://lcsb-databases.epfl.ch/atlas/.

## METHODS

In BridgIT, the Tanimoto score is used to quantify the similarity of reaction fingerprints. BridgIT allows us to do the following: (i) compare a given novel or orphan reaction to a set of reactions that have associated sequences, subsequently referred to as the *reference reactions*; (ii) rank the identified *similar* reactions based on the computed Tanimoto scores; and (iii) propose the sequences of the highest ranked *reference reactions* as possible candidates for encoding the enzyme of the given *de novo* or orphan reaction.

### Reactive site identification

An enzymatic reaction occurs when its substrate(s) fits into the binding site of an enzyme. Since the structure and geometry of the binding sites of enzymes are complex and most of the time not fully characterized, we proposed focusing on the similarity of the reactive sites of their substrates. Following this, we used the expert-curated, generalized reaction rules of BNICE.ch to identify the reactive sites of substrates. These reaction rules have third-level EC identifiers, e.g., EC 1.1.1, and they encompass the following biochemical knowledge of enzymatic reactions: (i) the information about atoms of the substrate’s reactive site; (ii) their connectivity (atom-bond-atom); and (iii) the exact information of bond breakage and formation during the reaction. As of July 2017, BNICE.ch contains 361 bidirectional generalized reaction rules that can reconstruct 6,528 KEGG reactions (44).

Given a novel or orphan reaction, the reactive sites of its substrate(s) are identified in three steps. In the first step, the BNICE.ch generalized reaction rules that can be applied to groups of atoms from the analyzed substrates are identified. Then, the information about the identified rules and the corresponding groups of atoms is stored. Subsequently, these groups of atoms are then referred to as the candidate substrate reactive sites. In the second step, among the identified rules, only the ones that can recognize the connectivity between the atoms of the candidate substrate reactive sites are kept. In the third step, whether the biotransformation of a substrate(s) to a product(s) can be explained by the rules retained after the second step is tested. The candidate reactive sites corresponding to the rules that have passed the three-step test are validated and used for the construction of reaction fingerprints.

We illustrate this procedure on the *de novo* reaction rat132064, which catalyzes the conversion of 3,4-dyhydroxymandelonitrile, substrate A, to protocatechualdehyde and cyanide (Fig. 1). In the first step, 164 rules were identified out of 361 rules that could be applied to groups of atoms of substrate A (Fig. 1, panel 1a). Out of the 164 rules, 103 matched the connectivity (Fig. 1, panel 1b). Finally, the 103 reaction rules were applied to substrate A for bond breaking and formation comparisons, and one rule could explain the transformation of substrate A to the products (Fig. 1, panel 1c).

### Reaction fingerprint construction

Molecular fingerprints, which are the linear representations of the structures of molecules, have been used in many methods and for different applications, especially for structural comparison of compounds (48, 49). One of the most commonly used molecular fingerprints is the Daylight fingerprint (48), and it decomposes a molecule into eight layers starting from layer zero that accounts only for atoms. Layer 1 expands one bond away from all of the atoms and accounts for atom-bond-atom connections. This procedure is continued until layer 7, which includes seven connected bonds from each atom. There are two types of Daylight reaction fingerprints: (i) structural reaction fingerprints, which are simple combinations of reactant and product fingerprints, and (ii) reaction difference fingerprints, which are the algebraic summation of reactant and product fingerprints multiplied by their stoichiometry coefficients in the reaction. In this study, we propose a modified version of the reaction difference fingerprint. The procedure for formulating BridgIT reaction fingerprints is demonstrated through an example reaction (Fig. 1, panel 2).

Starting from the atoms of the identified substrate reactive site, eight description layers of the molecule were formed, where different layers consisted of fragments with different lengths. Fragments were composed of atoms connected through unbranched sequences of bonds. Depending on the number of bonds included in the fragments, different description layers of a molecule were formed as follows:

Layer 0: Describes the type of each atom of the reactive site together with its count. For example, the substrate of the example reaction at layer 0 was described as 1 oxygen, 1 nitrogen, and 2 carbon atoms (Fig. 1, panel 2a).

Layer 1: Describes the type and count of each bond between pairs of atoms in the reactive site. In the example, the substrate at layer 1 was described with three fragments of length 1: 1 C-O, 1 C-C, and 1 C≡N bond (Fig. 1, panel 2a). Fragments are shown by their SMILES molecular representation (50).

Layer 2: Describes the type and count of fragments with three connected atoms. While layers 0 and 1 described the atoms of reactive sites, starting from layer 2, atoms that were outside of the reactive site were also described. In the illustrated example, there were three different fragments of this type (Fig. 1, panel 2a).

The same procedure was used to describe the molecules up to layer 7. Interestingly, and consistent with the previously reported result (41), we found that the 7-layer description was good enough to capture the structure of most of the metabolites in biochemical reactions, therefore providing a precise reaction fingerprint. Note that not all description layers are needed to describe less complex molecules. For example, product C (cyanide) was fully described using only layer 0 and layer 1 (Fig. 1, panel 2a). For very large molecules, the description layers that contain fragments with more than 8 connected atoms can be used.

For each layer, the substrate set was formed by merging all of the fragments, their type, and their count in the substrate molecules of the reaction, and the product set was formed by merging all of the fragments (type and count) in the product molecules of the reaction. In both sets, the count of each fragment was multiplied by the stoichiometric coefficients of the corresponding compound in the reaction. Finally, the reaction fingerprints were created by summing the fragments of the substrate and product sets for each layer (Fig. 1, panel 2b).

Introducing the specificity of reactive sites into the reaction fingerprint allows BridgIT to capitalize on the information about enzyme binding pockets (15). To keep this valuable information throughout the generation of reaction fingerprints, BridgIT does not consider the atoms of the reactive site(s) when performing the algebraic summation of the substrate and product set fragments. Consequently, the BridgIT algorithm enables retaining, tracking, and emphasizing the information of the reactive site(s) in all of the layers of the reaction fingerprint, which distinguishes it from the existing methods.

### Reaction similarity evaluation

The similarity of two reactions was quantified using the similarity score between their fingerprints, subsequently referred to as reaction fingerprints A and B. In this study, the Tanimoto score, which is an extended version of the Jaccard coefficient and cosine similarity, was used (51). Values of the Tanimoto scores near 0 indicate reactions with no or negligible similarity, whereas values near 1 indicate reactions with high similarity.

The Tanimoto score for each descriptive layer, *T_LK_*, together with the global Tanimoto score, *T_G_*, was calculated. The Tanimoto score for the k-th descriptive layer was defined as:

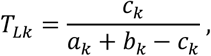

where *a_k_* was the count of the fragments in the k-th layer of reaction fingerprint A; *b_k_* was the count of the fragments in the k-th layer of reaction fingerprint B; and *c_k_* was the number of common k-th layer fragments of reaction fingerprints A and B. Two fragments were equal if their canonical SMILES and their stoichiometric coefficients were identical. The global Tanimoto similarity score, *T_G_*, was defined as follows:

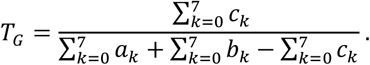

For each reaction fingerprint, its Tanimoto similarity score was calculated against the reaction fingerprints from the BridgIT reference database, which contained reaction fingerprints of all known, well-characterized enzymatic reactions (Fig. 1, panel 3).

### Sorting, ranking and gene assignment

For a given input reaction, the reference reactions were ranked using the computed T_G_ scores. The algorithm distinguished between the identified reference reactions with the same T_G_ score based on the T_L_ score of layers 0 and 1, and it also allows the user to assign ranking weights to specified layers. The protein sequences associated with the highest ranked, i.e., the most similar, reference reactions were then assigned to the input reaction (Fig. 1, panel 4).

## CONCLUSIONS

We developed the computational tool, BridgIT, to evaluate and quantify the structural similarity of biochemical reactions by exploiting the biochemical knowledge of BNICE.ch generalized reaction rules. Because the generalized reaction rules can identify reactive sites of substrates, BridgIT can translate the structural definition of biochemical reactions into a novel type of reaction fingerprint that explicitly describes the atoms of the substrates’ reactive sites and their surrounding structure. Through the analysis of 5,049 known and well-defined biochemical reactions, we found that knowledge of the neighborhood up to three bonds away from the atoms of the reactive site can predict biochemistry and match catalytic protein sequences. The reaction fingerprints proposed in this work can be used to compare all novel and orphan reactions to well-characterized reference reactions and, consequently, to link them with genes, genomes, and organisms. We demonstrated through several examples the improvements that the BridgIT fingerprint brings to the field compared to the fingerprints currently existing in the literature.

A drawback of traditional sequence similarity methods is that they cannot identify protein sequence candidates for *de novo* reactions, which we have shown BridgIT can do. We tested BridgIT predictions against experimental biochemical evidence, within two large-scale validations studies on sets of (i) 234 orphan and (ii) 379 *de novo* reactions. The reactions from these two sets were unknown in the previous versions of the KEGG database but were later experimentally confirmed and catalogued in KEGG 2018. BridgIT predicted the exact or a highly related enzyme for 89% of these reactions.

We further applied BridgIT to the entire catalog of *de novo* reactions of the ATLAS of Biochemistry database and proposed several candidate enzymes for each of them. The candidate enzymes for these *de novo* reactions are either immediately capable of catalyzing these reactions or can serve as initial sequences for enzyme engineering. The obtained BridgIT similarity scores can also be used as a confidence score to assess the feasibility of the implementation of novel ATLAS reactions in metabolic engineering and systems biology studies.

The applications of BridgIT go beyond merely bridging gaps in metabolic reconstructions, as this method can be used to identify the potential utility of existing enzymes for bioremediation as well as for various applications in synthetic biology and metabolic engineering. As the field of metabolic engineering grows and metabolic engineering applications increasingly turn towards the production of valuable industrial chemicals such as 1,4-butanediol (52, 53), we expect that methods for the design of *de novo* synthetic pathways, such as BNICE.ch (15), and methods for identifying candidate enzymes for *de novo* reactions, such as BridgIT, will grow in importance.

## Abbreviations

KEGG: Kyoto Encyclopedia of Genes and Genomes
RF: Reaction fingerprint
DT: Discrimination Threshold
TP: True Positives
TN: True Negatives
FP: False Positives
FN: False Negatives
ROC: Receiver Operating Characteristics
AUC: Area Under a ROC Curve

## REFERENCES

1. Orth JD, Conrad TM, Na J, Lerman JA, Nam H, Feist AM, et al. A comprehensive genome-scale reconstruction of Escherichia coli metabolism—2011. Mol Syst Biol. 2011 Jan 1;7(1):535.

2. Kanehisa M, Furumichi M, Tanabe M, Sato Y, Morishima K. KEGG: new perspectives on genomes, pathways, diseases and drugs. Nucleic Acids Res. 2017 Jan 4;45(D1):D353–61.

3. Sorokina M, Stam M, Medigue C, Lespinet O, Vallenet D. Profiling the orphan enzymes. Biol Direct. 2014;9.

4. Gao J, Ellis LBM, Wackett LP. The University of Minnesota Biocatalysis/Biodegradation Database: improving public access. Nucleic Acids Res. 2010 Jan;38(suppl_1):D488–91.

5. Hatzimanikatis V, Li CH, Ionita JA, Henry CS. Exploring the diversity of complex metabolic networks. Bioinformatics. 2005;21:1603–1609.

6. Hatzimanikatis V, Li C, Ionita JA, Broadbelt LJ. Metabolic networks: enzyme function and metabolite structure. Curr Opin Struct Biol. 2004 Jun;14(3):300–6.

7. Soh KC, Hatzimanikatis V. DREAMS of metabolism. Trends Biotechnol. 2010 Oct;28(10):501–8.

8. Carbonell P, Planson A-G, Fichera D, Faulon J-L. A retrosynthetic biology approach to metabolic pathway design for therapeutic production. BMC Syst Biol. 2011;5(1):122.

9. Rodrigo G, Carrera J, Prather KJ, Jaramillo A. DESHARKY: automatic design of metabolic pathways for optimal cell growth. Bioinformatics. 2008 Nov 1;24(21):2554–6.

10. Cho A, Yun H, Park J, Lee S, Park S. Prediction of novel synthetic pathways for the production of desired chemicals. BMC Syst Biol. 2010;4(1):35.

11. Yim H, Haselbeck R, Niu W, Pujol-Baxley C. Metabolic engineering of Escherichia coli for direct production of 1,4-butanediol. Nat Chem Biol. 2011;445–452.

12. Campodonico MA, Andrews BA, Asenjo JA, Palsson BO, Feist AM. Generation of an atlas for commodity chemical production in Escherichia coli and a novel pathway prediction algorithm, GEM-Path. Metab Eng. 2014;25:140–158.

13. Prather KLJ, Martin CH. De novo biosynthetic pathways: rational design of microbial chemical factories. Curr Opin Biotechnol. 2008 Oct;19(5):468–74.

14. Delépine B, Duigou T, Carbonell P, Faulon J-L. RetroPath2.0: A retrosynthesis workflow for metabolic engineers. 2017 Jun 29 [cited 2017 Aug 18]; Available from: http://biorxiv.org/lookup/doi/10.1101/141721

15. Hadadi N, Hatzimanikatis V. Design of computational retrobiosynthesis tools for the design of de novo synthetic pathways. Curr Opin Chem Biol. 2015;28:99–104.

16. Carbonell P, Parutto P, Herisson J, Pandit SB, Faulon JL. XTMS: pathway design in an eXTended metabolic space. Nucleic Acids Res. 2014;42:389–394.

17. Hadadi N, Soh KC, Seijo M, Zisaki A. A computational framework for integration of lipidomics data into metabolic pathways. Metab Eng. 2014;23:1–8.

18. Rolfsson O, Palsson BØ, Thiele I. The human metabolic reconstruction Recon 1 directs hypotheses of novel human metabolic functions. BMC Syst Biol. 2011;5(1):155.

19. Karp PD. Call for an enzyme genomics initiative. Genome Biol. 2004;5.

20. Orth JD, Palsson BO. Systematizing the Generation of Missing Metabolic Knowledge. Biotechnol Bioeng. 2010;107:403–412.

21. Osterman A, Overbeek R. Missing genes in metabolic pathways: a comparative genomics approach. Curr Opin Chem Biol. 2003;7:238–251.

22. Overbeek R, Fonstein M, D’Souza M, Pusch GD, Maltsev N. The use of gene clusters to infer functional coupling. In: Proceedings of the National Academy of Sciences of the United States of America 1999, 96. p. 2896–2901.

23. Pellegrini M, Marcotte EM, Thompson MJ, Eisenberg D, Yeates TO. Assigning protein functions by comparative genome analysis: Protein phylogenetic profiles. In: Proceedings of the National Academy of Sciences of the United States of America 1999, 96. p. 4285–4288.

24. Chen V. Predicting genes for orphan metabolic activities using phylogenetic profiles. Genome Biol. 2006;17.

25. Overbeek R, Begley T, Butler RM, Choudhuri JV. The subsystems approach to genome annotation and its use in the project to annotate 1000 genomes. Nucleic Acids Res. 2005;33:5691–5702.

26. Vallenet D, Labarre L, Rouy Z, Barbe V. a microbial genome annotation system supported by synteny results. Nucleic Acids Res. 2006;34:53–65.

27. Kharchenko P, Chen LF, Freund Y, Vitkup D, Church GM. Identifying metabolic enzymes with multiple types of association evidence. BMC Bioinformatics. 2006; 7.

28. Yamanishi Y, Mihara H, Osaki M, Muramatsu H. Prediction of missing enzyme genes in a bacterial metabolic network - Reconstruction of the lysine-degradation pathway of Pseudomonas aeruginosa. Febs J. 2007;274:2262–2273.

29. Chen Y, Mao FL, Li G, Xu Y. Genome-wide discovery of missing genes in biological pathways of prokaryotes. BMC Bioinformatics. 2011;12.

30. Smith AAT, Belda E, Viari A, Medigue C, Vallenet D. The CanOE Strategy: Integrating Genomic and Metabolic Contexts across Multiple Prokaryote Genomes to Find Candidate Genes for Orphan Enzymes. Plos Comput Biol. 2012;8.

31. Schnoes AM, Brown SD, Dodevski I, Babbitt PC. Annotation Error in Public Databases: Misannotation of Molecular Function in Enzyme Superfamilies. Plos Comput Biol. 2009;5.

32. Green ML, Karp PD. Using genome-context data to identify specific types of functional associations in pathway/genome databases. Bioinformatics. 2007;23:205–211.

33. Matsuta Y, Ito M, Tohsato Y. ECOH: An Enzyme Commission number predictor using mutual information and a support vector machine. Bioinformatics. 2013;29:365–372.

34. Galperin MY, Koonin EV. Divergence and Convergence in Enzyme Evolution. J Biol Chem. 2012 Jan 2;287(1):21–8.

35. Ofran Y, Margalit H. Proteins of the same fold and unrelated sequences have similar amino acid composition. Proteins Struct Funct Bioinforma. 2006 Jul 1;64(1):275–9.

36. Giri V, Sivakumar TV, Cho KM, Kim TY, Bhaduri A. RxnSim: a tool to compare biochemical reactions. Bioinformatics. 2015;31:3712–3714.

37. Hu QN, Deng Z, Hu HA, Cao DS, Liang YZ. RxnFinder: biochemical reaction search engines using molecular structures, molecular fragments and reaction similarity. Bioinformatics. 2011;27:2465–2467.

38. Moriya Y, Yamada T, Okuda S, Nakagawa Z. Identification of Enzyme Genes Using Chemical Structure Alignments of Substrate-Product Pairs. J Chem Inf Model. 2016;56:510–516.

39. Hu QN, Zhu H, Li XB, Zhang MM. Assignment of EC Numbers to Enzymatic Reactions with Reaction Difference Fingerprints. Plos One. 2012;7.

40. Rahman SA, Cuesta SM, Furnham N, Holliday GL, Thornton JM. EC-BLAST: a tool to automatically search and compare enzyme reactions. Nat Methods. 2014 Feb;11(2):171–4.

41. DAYLIGHT, Version 4.62, DAYLIGHT Inc., Mission Viejo, CA.

42. Rogers DJ, Tanimoto TT. A Computer Program for Classifying Plants. Science. 1960(132):1115–1118.

43. International Union of Biochemistry and Molecular Biology, Webb EC, editors. Enzyme nomenclature 1992: recommendations of the Nomenclature Committee of the International Union of Biochemistry and Molecular Biology on the nomenclature and classification of enzymes. San Diego: Published for the International Union of Biochemistry and Molecular Biology by Academic Press; 1992. 862 p.

44. Hadadi N, Hafner J, Shajkofci A, Zisaki A, Hatzimanikatis V. ATLAS of Biochemistry: A Repository of All Possible Biochemical Reactions for Synthetic Biology and Metabolic Engineering Studies. ACS Synth Biol. 2016 Oct 21;5(10):1155–66.

45. Hadadi N, Hafner J, Soh KC, Hatzimanikatis V. Reconstruction of biological pathways and metabolic networks from in silico labeled metabolites. Biotechnol J. 2017 Jan;12(1):1600464.

46. Pundir S, Magrane M, Martin MJ, O’Donovan C, The UniProt Consortium. Searching and Navigating UniProt Databases: Searching and Navigating UniProt Databases. In: Bateman A, Pearson WR, Stein LD, Stormo GD, Yates JR, editors. Current Protocols in Bioinformatics [Internet]. Hoboken, NJ, USA: John Wiley & Sons, Inc.; 2015 [cited 2017 Aug 18]. p. 1.27.1–1.27.10. Available from: http://doi.wiley.com/10.1002/0471250953.bi0127s50

47. Altschul SF, Gish W, Miller W, Myers EW, Lipman DJ. Basic Local Alignment Search Tool. J Mol Biol. 1990;215:403–410.

48. Briem H, Lessel UF. In vitro and in silico affinity fingerprints: Finding similarities beyond structural classes. In: Klebe G, editor. Virtual Screening: An Alternative or Complement to High Throughput Screening? [Internet]. Dordrecht: Kluwer Academic Publishers; 2002 [cited 2017 Aug 18]. p. 231–44. Available from: http://link.springer.com/10.1007/0-306-46883-2_13

49. O’Boyle NM, Banck M, James CA, Morley C, Vandermeersch T, Hutchison GR. Open Babel: An open chemical toolbox. J Cheminformatics. 2011;3(1):33.

50. Weininger D. SMILES, a chemical language and information system. 1. Introduction to methodology and encoding rules. J Chem Inf Comput Sci. 1988 Feb 1;28(1):31–6.

51. Leydesdorff L. On the normalization and visualization of author co-citation data: Salton’s Cosineversus the Jaccard index. J Am Soc Inf Sci Technol. 2008 Jan 1;59(1):77–85.

52. Burgard A, Burk MJ, Osterhout R, Van Dien S, Yim H. Development of a commercial scale process for production of 1,4-butanediol from sugar. Curr Opin Biotechnol. 2016;42:118–125.

53. Andreozzi S, Chakrabarti A, Soh KC, Burgard A. Identification of metabolic engineering targets for the enhancement of 1,4-butanediol production in recombinant E. coli using large-scale kinetic models. Metab Eng. 2016;35:148–159.

